# CRISPR-induced deletion with SaCas9 restores dystrophin expression in dystrophic models *in vitro* and *in vivo*

**DOI:** 10.1101/378331

**Authors:** Benjamin L. Duchêne, Khadija Cherif, Jean-Paul Iyombe-Engembe, Antoine Guyon, Joel Rousseau, Dominique L. Ouellet, Xavier Barbeau, Patrick Lague, Jacques P. Tremblay

## Abstract

Duchenne Muscular Dystrophy (DMD), a severe hereditary disease, affecting 1 boy out of 3500, mainly results from the deletion of one or more exons leading to a reading frame shift of the *DMD* gene that abrogates dystrophin protein synthesis. We used the Cas9 of *Staphylococcus aureus* (SaCas9) to edit the human *DMD* gene. Pairs of sgRNAs were meticulously chosen to induce a genomic deletion to not only restore the reading frame but also produced a dystrophin protein with normally phased spectrin-like repeats. The formation of a dystrophin protein with spectrin-like repeats normally phased is not usually obtained by skipping or by deletion of complete exons. This can however be obtained in rare instances where the exon/intron borders of the beginning and the end of the complete deletion (patient deletion plus CRISPR-induced deletion are at similar positions in the spectrin-like repeat. We used pairs of sgRNAs, targeting exons 47 and 58 and a normal reading frame was restored in 67 to 86% of the resulting hybrid exons in myoblasts derived from muscle biopsies of 4 DMD patients with different exon deletions. The restoration of the DMD reading frame and restoration of the dystrophin expression was also obtained *in vivo* in the heart of the del52hDMD/mđx. Our results provide a proof-of-principle that SaCas9 could be used to edit the human DMD gene and could be considered for the further development of a therapy for DMD.

## INTRODUCTION

Duchenne muscular dystrophy is one of the most severe hereditary disease affecting 1 in 3500 new-born boys^1, 2^ This X-linked disorder is caused by a mutation in the *DMD* gene coding for the dystrophin protein leading to the absence of this protein in muscle fibers^3, 4^ These mutations, mostly deletions of one or several exons, are responsible of a shift in the normal reading frame of the *DMD* gene thus generating a premature stop codon leading to the loss of the dystrophin protein during the translation process. This protein is fundamental to maintain the integrity of the sarcolemma during contraction as it allows interaction with the surrounding extracellular matrix through binding with the membrane-anchored β-dystroglycan and with the cytoskeleton through interaction with actin and microtubules^5^. Notably, the dystrophin central rod domain is formed by 24 successive spectrin-like repeats (SLRs). Each of these SLRs is formed by three antiparallel α-helixes (A, B and C) forming a coiled-coiled structure, in which each heptad contains hydrophobic amino acids in positions “a” and “d”. Interestingly, the nNOS interacts with the helixes A, B and C of the SLR 16 and with the helixes A of the SLR 17. It has been reported that in Becker muscular dystrophy (BMD) patients^6^, the severity of the disease could be related to the structure of their truncated dystrophin, thus we focused on creating a genomic deletion that not only restores the *DMD* gene reading frame but also allows the production of a dystrophin protein with SLRs correctly phased and structured. Recent advances in the field of gene therapy offers great perspectives for the development of a curative treatment for DMD. Strategies of gene replacement that rely on the AAV-mediated delivery of micro-dystrophin still undergo development^7–9^ and reached clinical trial phase such as the rAAVrh74.MHCK7.micro-dystrophin (Nationwide Children’s Hospital Columbus, Ohio, United States) or the SGT-001 (University of Florida, Gainesville, Florida, United States) (https://clinicaltrials.gov/). However, few years ago, the discovery of the CRISPR/Cas9 system raised hope for the establishment of therapies for hereditary diseases, that counter the disease at its root, by editing the genome. This system was first identified as the immune system of some bacteria species such as *Streptococcus pyogenes* or *Staphylococcus aureus*, to protect them against bacteriophage infections^10^. This system has been adapted to target genomic DNA in mammals, thus offering possibilities to permanently edit their genome^11–13^ More precisely, a single guide RNA (sgRNA) can be made to target a specific protospacer sequence as long as it is positioned next to a protospacer adjacent motif (PAM), which is NGG for S. *pyogenes* and NNGRRT for S. *aureus.* Once the Cas9/sgRNA complex binds to its protospacer, Cas9 protein is able to generate a double strand break precisely three nucleotides upstream of the SpCas9 or the SaCas9 PAM. Based on this precise editing capacity, we have previously shown that by using a combination of two sgRNAs, we can generate a genomic deletion that forms a hybrid exon 50-54, which restores the dystrophin protein expression in myotubes from a patient having a deletion of exons 51 to 53^14, 15^ In DMD patients, reported mutations are mostly comprised in the “hot spot” including exons 45 to 55. To date, most of the research groups focus on the use of the CRISPR/Cas9 system to restore a normal reading-frame through the complete removal of one or several exons. which is not taking care to restore the normal phasing of spectrin-like repeat. Such approach could lead to the formation of a dystrophin protein with an abnormally structured central rod-domain, which can lead to a severe Becker Muscular Dystrophy instead of a milder BMD. Here we report an approach that covers up to 40% of all *DMD* mutations by creating a hybrid exon directly connecting exons 47 to 58 using the SaCas9. Please note that we did not form a hybrid exon 44-58 nor 45-58 because the dystrophin sequences of the SLR R17, coded by exons 44 to 46, are necessary for the essential interaction of dystrophin with nNOS^16^. Our approach will not only restore the dystrophin production but will also produce a dystrophin protein with a normal SLR structure. More precisely, by using a combination of 2 sgRNAs, we created a hybrid exon allowing to position hydrophobic amino acids in positions “a” and “d” of the newly formed SLR. We think that by maintaining the adequate structure of the central rod domain, i.e., the SLRs, we can not only maintain the interaction of dystrophin with elements of the dystrophin-associated proteins and with nNOS but also insure the normal function of the dystrophin protein during contraction and relaxation of the muscle fibres. To precisely produce a normally phased SLR, we used our previously characterized CRISPR-induced deletion method (CinDel method) that aims to target into the exons that precede and follow the patient mutation to form a hybrid exon. We have, however, modified this approach to use the smaller SaCas9 from S. *aureus* whose gene could ultimately be delivered with a pair of sgRNAs by a single AAV^17^ Besides, we are not targeting the exons that immediately precede and follow the patient deletion but rather the exons 47 and 58 so that the formation of the hybrid exon 47-58 may be a therapeutic approach for any deletion, insertion or point mutations between these two exons. Our proposed edition would thus be a treatment for roughly 40% of DMD patients.

## RESULTS

### Identification and activity assays of individual sgRNAs in 293T cells

Our primary goal was to establish a strategy based on the creation of a hybrid exon allowing the correction of the *DMD* gene reading frame, in the case of DMD patients affected by the deletion of exon 50. Thus, all possible sgRNAs target sites in exons 46 to 58 were screened. We screened 51 sgRNA target sites for the SaCas9 nuclease. Since the SaCas9 nuclease induces a double strand break (DSB) at precisely 3 nucleotides upstream of the PAM (NNGRRT), we were able to select pairs of sgRNAs that can be combined to create a new hybrid exon that restores a normal reading frame. Thus, these hybrid exons might permit the production of an internally deleted dystrophin protein. In addition, we completed our analysis by selecting pairs of sgRNAs that were also able to produce an adequate SLR. Consequently, we focused only on combinations where the hybrid junction maintained the configuration of a normal SLR where hydrophobic amino acids are localized in position “a” and “d” of the heptad motif (SLR are composed of 3 α-helixes, each containing 7 amino-acids (a to f) where hydrophobic amino acid are in the location “a” and “d”). In order to assess the adequate localization of those amino acids, we referred to the eDystrophin database, which provides information about the structural domains of the dystrophin protein (http://edystrophin.genouest.org/). As a result of our preliminary analysis, we focused our work on 18 sgRNAs (**table 1 and Supplementary Figures S1A & B**), which can in principle generate 12 hybrid exons that do not only correct the reading frame of the *DMD* gene but also maintained the structure of the SLRs in the central rod domain (**Supplementary Figure S1C**).

**Table 1.**
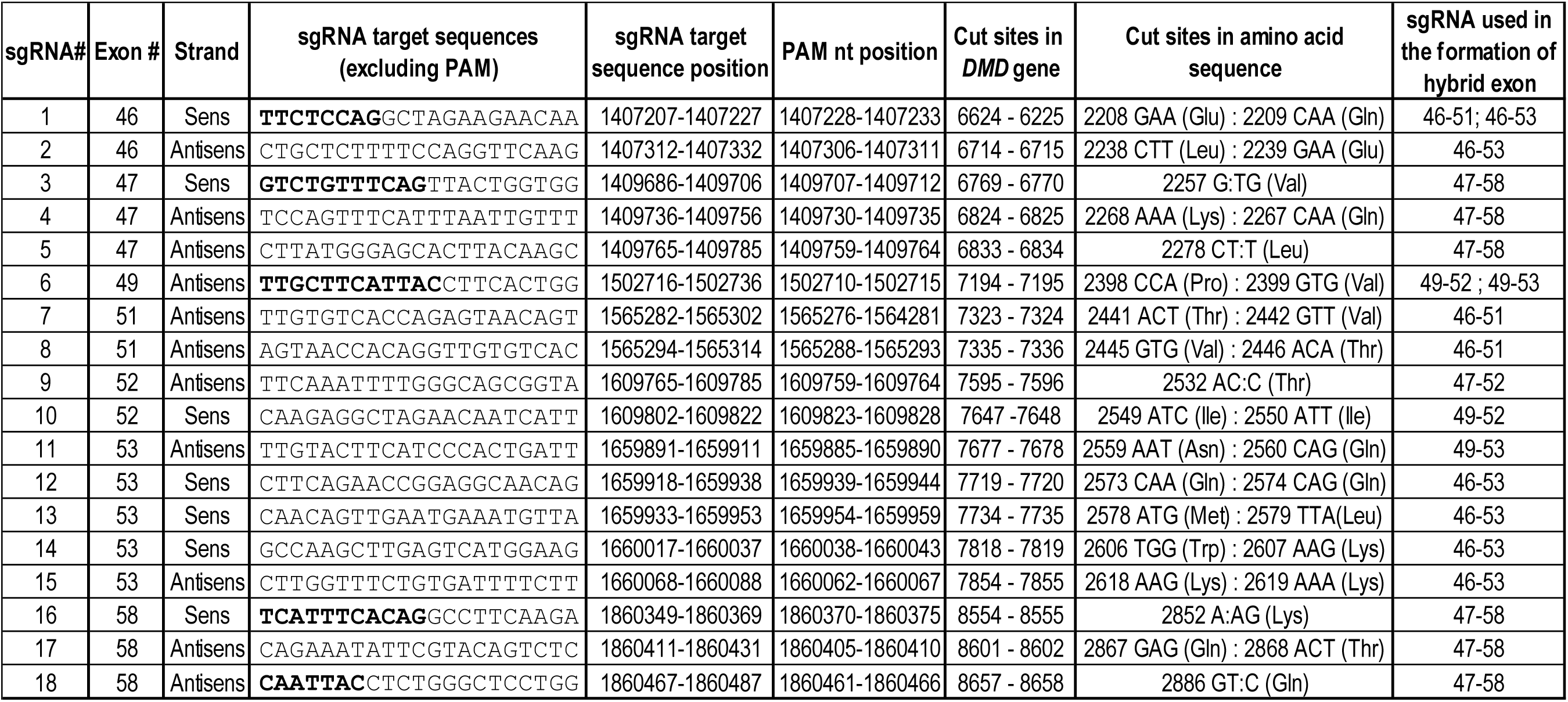
List of sgRNAs target sites in exons 46 to 58. Nucleotides position are provided with reference to the *DMD* gene sequence ENS00000198947 analysed using Benchling web tool (https://benchling.com/).

Next, we designed 18 sgRNAs, which were cloned into the PX601-AAV-CMV::NLS-SaCas9-NLS-3xHA-bGHpA;U6::BsaI-sgRNA plasmid (Addgene # 61591). These plasmids were, first individually, transfected in 293T cells to assess their activity. Genomic DNA was extracted 48 hours post transfection. The targeted exon, either 46, 47, 49, 51, 52, 53 or 58 was amplified by PCR and submitted to a Surveyor assay for the detection of INDELs (**Supplementary Figure S2A**). Among the 18 sgRNAs tested, sgRNA 2-46, sgRNA 4-47, sgRNA 9-52, sgRNA 11-53, sgRNA 15-53 showed little or no activity whereas all remaining sgRNAs exhibited good cleavage efficiency as observed with the Surveyor assays that generated cleaved bands at the expected sizes. To improve the characterization of the formation of INDELs at the targeted sites, PCR products were submitted to TIDE analysis. The percentages of INDELs determined by this method were consistent with the Surveyor assay as samples that exhibited a substantial cleavage activity also demonstrated a high rate of INDELs according to the TIDE analysis (**Supplementary Figure S2B**).

### Test of sgRNA pairs in 293T cells

293T cells were co-transfected with pairs of sgRNAs that might not only produce a large genomic deletion but also precisely connect exons, surrounding the deletion of the exon 50, to restore the reading frame and produce an adequate SLR. Thus, 12 pairs of sgRNAs (previously identified in a preliminary analysis of compatible sgRNAs) were tested. To detect the formation of the hybrid exons, PCRs were performed using the forward primer binding with the targeted exon upstream of the exon 50 and the reverse primer binding with the targeted exon downstream of the exon 50, previously used when sgRNAs were individually tested (**Figure 1**). Given that the targeted exons are separated by large introns, PCR amplification was only possible if exon deletion had occurred. The combination of sgRNAs 2-46 and 15-53 did not allow the formation a hybrid exon 46-53 thus no PCR product was detected. In addition, sgRNA 5-47 combined with sgRNA 9-52 and sgRNA 4-47 combined with sgRNA 17-58 were poorly effective for the formation of the hybrid exons 47-52 and 47-58 respectively. However, all the remaining combinations of sgRNAs permitted the formation of hybrid exons as observed by the detection of PCR products in these conditions.

**Figure 1.**
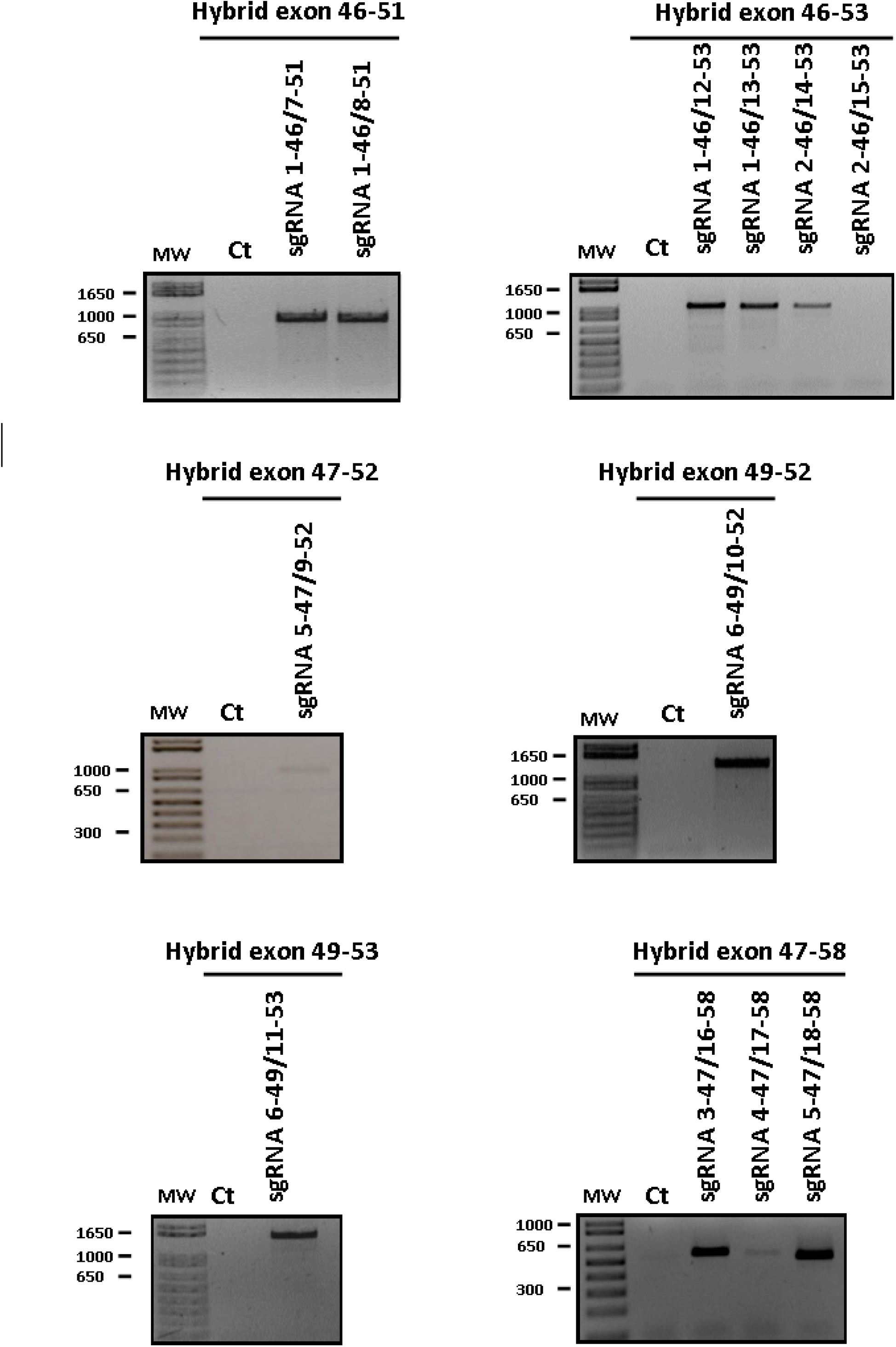
Generation of several hybrid exons by genomic edition using pairs of sgRNAs. (A) The formation of hybrid exons was tested with various pairs of sgRNAs, previously characterized in Table 2, which could in principle restore an open reading-frame for the *DMD* gene. The formation of the hybrid exon was confirmed by PCR amplification, (i.e., there was no PCR product when the hybrid exon is not formed). As a negative control (Ct) we used non-transfected 293T cells.

### Analysis of hybrid exons

Before proceeding to further analyses, we needed to confirm that the hybrid exons that were formed were correct. Indeed, these hybrid exons should predominantly be able to restore a normal reading frame for the *DMD* gene. The PCR products resulting from the amplification of hybrid exons were purified and cloned into a pMiniT vector. For the analysis of the sequencing results of the hybrid exons, we counted the bacterial clones that contained the right nucleotide sequence and the bacterial clones that contained a hybrid exon where the nucleotide sequence was not exactly as expected (data not shown). To improve the characterization of these hybrid exons, we performed a modified TIDE analysis that compared a PCR product composed of 100% of the expected hybrid exon (amplified from a previously identified clone that contained a hybrid exon with the right nucleotide sequence), against a test sample amplified from the genomic DNA of treated cells (see **Table 2**). In at least 54% of the PCR products, the nucleotide sequences of the hybrid exons were exactly as expected (**Supplementary data - Figure S3**), and thus efficiently restored a normal reading frame in the *DMD* gene. Interestingly, the combination of sgRNAs 3-47 and 16-58, which resulted in a 450 kbp deletion was one of the most precise pairs since 90% of the PCR products of the resulting hybrid exon 3-47/16-58 were exactly as expected. In addition, the combination of sgRNAs 5-47 and 18-58 also generated an interesting large genomic deletion, although only 68% of the PCR products contained the expected nucleotide sequence. The formation of the hybrid exons 47-58 could be a treatment for about 40% of the mutations (deletion, duplications and point mutations) identified in DMD patients. Consequently, in the next steps of our work, we choose to focus on these two combinations only.

**Table 2.** Analysis of hybrid exon amplicons by TIDE.

| Upstream<br>sgRNA | Downstream<br>sgRNA | Expected size of<br>the amplicon (bp) | Percentage of<br>correct junction |
| --- | --- | --- | --- |
| 1-46 | 7-51 | 967 | 90,8 |
| 1-46 | 8-51 | 955 | 86,1 |
| 1-46 | 12-53 | 1099 | 54,6 |
| 1-46 | 13-53 | 1084 | 60,9 |
| 2-46 | 14-53 | 1090 | N/A |
| 2-46 | 15-53 | 1054 | N/A |
| 5-47 | 9-52 | 997 | N/A |
| 6-49 | 10-52 | 1386 | 83,3 |
| 6-49 | 11-53 | 1548 | 84,7 |
| 3-47 | 16-58 | 587 | 90,7 |
| 4-47 | 17-58 | 575 | N/A |
| 5-47 | 18-58 | 548 | 67,7 |

### 3D spectrin-like models resulting from the formation of hybrid exons 3—47/16—58 and 5—47/18—58

The dystrophin protein is important to maintain the integrity of the sarcolemma. The central part of this protein, called the rod domain, is made of 24 SLRs. According to homology models available from the eDystrophin Website (http://edystrophin.genouest.org/), the dystrophin spectrin-like repeat predicted structures are composed of the typical triple coiled-coil structure, i.e., three α-helices (A, B, and C) linked by two loops (AB and BC)^18^, as represented in **figure 2B** for the integral structure of the spectrin-like repeat 18 (R18). For the structure of the resulting hybrid spectrin-like repeat (R18-R23) using pairs of sgRNAs (either pair 3–47/16–58 or pair 5–47/18–5), the spectrin repeats R19 to R22 are removed but the reading frame resulting in the formation of hybrid exons 47–58 are conserved, both with junction points in Helix B (**Figure 2C & 2D**). The homology model for these two hybrid exons presents similar length and structure to the original spectrin-like repeat R18, with a shorter BC loop for hybrid exon 5–47/18–58. Thus, the homology models suggest that the dystrophin produced following the formation of hybrid exons with sgRNA pairs 3–47/16–58 or 5–47/18–58 would have a similar structure as the integral dystrophin and therefore should function in a similar manner. These predictive structures of hybrid SLRs confirmed that the hybrid exons 47–58 (formed with either with sgRNAs 3–47 and 16–58 or with sgRNAs 5–47 and 18–58) are good candidates to produce dystrophin proteins with an adequate structure thus increasing the potential functionality of the newly produced internally-deleted dystrophin.

**Figure 2:**
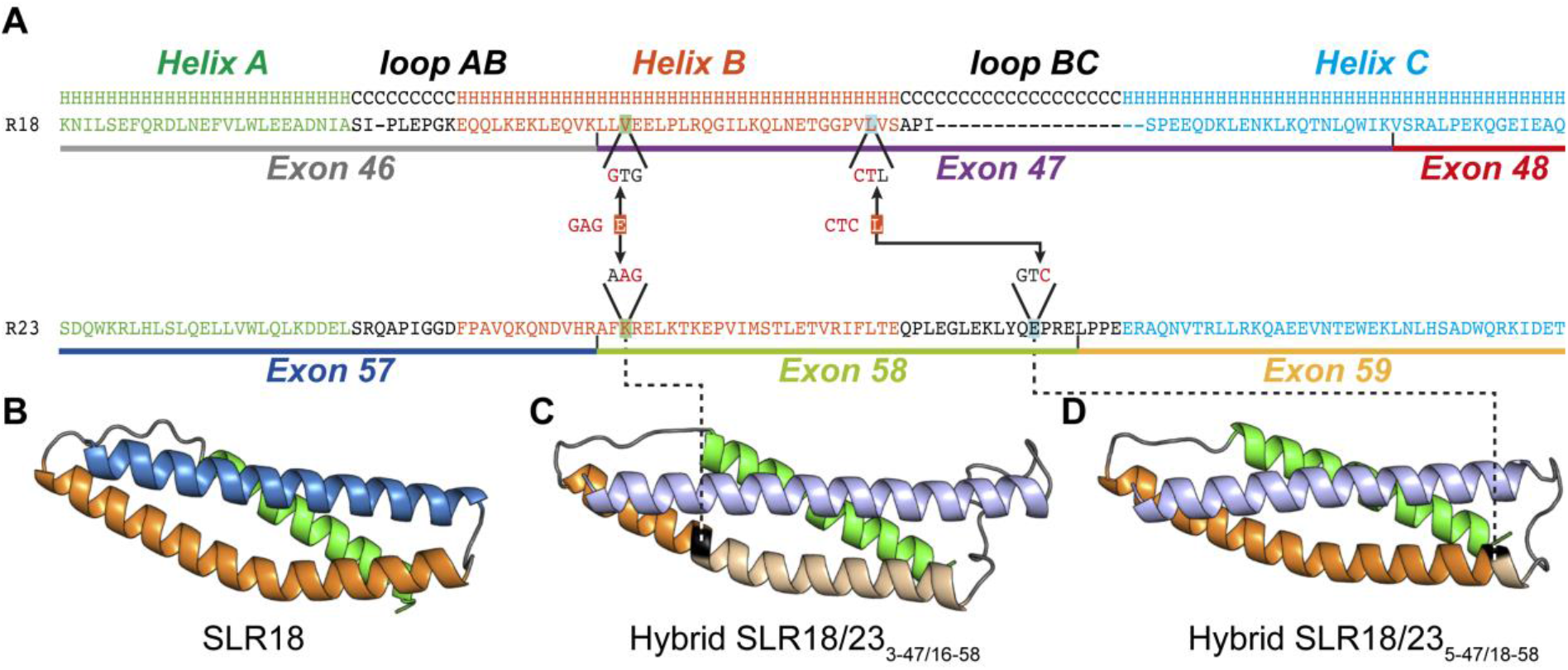
Structural representations of SLR18 and two hybrid SLRs. (A) Primary structure alignments for SLR18 and SLR23. The amino acid sequences of the exons associated with these spectrin-like repeats are underlined in different colours. The secondary structure of the SLRs is represented above the sequences, H for helices and C for the loop segments. Structural motifs, as identified in the primary sequence alignment, are coloured as follows: Helix A is in green, Helix B is in orange, and Helix C is in blue. Loops AB and BC are in black. The sgRNAs 3-47 and 5-47 are cutting in exon 47 in sequences coding for Helix B. The sgRNAs 16-58 and 18-58 are cutting respectively in exon 58 in sequences coding respectively for the Helix B and the BC loop. When these sgRNAs are used in pairs (i.e., sgRNAs 3-47 with 16-58 and sgRNAs 5-47 and 18-58), the nucleotides between their cut sites are deleted leading in both cases to the formation of the hybrid exon 47-58. Note that since these sgRNAs are cutting within a codon, a new codon is formed at the junction site, one coding for glutamic acid (E) and the other for leucine (L), respectively within hybrid exon 3-47/16-58 and 5-47/18-58. These hybrid exons produced respectively Hybrid SLR 3-47/16-58 and Hybrid SLR 5-47/18-58. (B) Homology models for integral spectrin-like repeat SLR18 obtained from eDystrophin Website (http://edystrophin.genouest.org/). (C) The homology model for the hybrid SLR18/23_3-47/16-58_ and (D) hybrid SLR18/23_5-47/18-58_ were realized using the iTasser web server (http://zhanglab.ccmb.med.umich.edu/I-TASSER/). Colours are darker for the segment derived from SLR18 and lighter for the segment derived from SLR23. Junction position is highlighted in black.

### Formation of the hybrid exons 47-58 and restoration of dystrophin protein expression in DMD patient cells

For the experiments with DMD patient cells, we only focused on the two pairs of sgRNAs that could be suitable for the highest number of *DMD* mutations, i.e., the combination of sgRNAs 3-47 and 16-58 and the combination of sgRNAs 5-47 and 18-58. First, transfection and nucleofection of a combination of plasmids that code for the SaCas9 and for a pair of sgRNAs were tested (data not shown). Due to low transfection and nucleofection efficiencies, viral transduction of myoblasts was subsequently tested. Using integrative lentiviral vectors encoding the SaCas9 gene and a pair of sgRNAs (i.e. Lenti-SaCas9-3-47/16-58 and Lenti-SaCas9-5-47/18-58) and eGFP, myoblasts from four different DMD patients having different exon deletions (Δ49-50, Δ50-52, Δ51-53 and Δ51-56) were transduced. The transduction was evaluated by eGFP expression in myoblasts three days later (**Figure 3A**). Following proliferation of transduced cells, genomic DNA was extracted from a cell sample to assess the formation of the hybrid exons 47-58 (**Figure 3B**). In non-transduced myoblasts, no PCR amplification was detected because the distance between the primers is too long (i.e., 450 kbp). In all transduced cells, the hybrid exons were detected with both pairs of sgRNAs (i.e., sgRNAs 3-47 and 16-58, and sgRNAs 5-47 and 18-58). In addition, PCR products were submitted to sequencing to confirm that the hybrid exons contained the expected nucleotide sequences. We observed by TIDE analysis, that the hybrid exons contained the exact nucleotide sequences in 67% and 73%, respectively for the pairs of sgRNAs 3-47/16-58 and 5-47/18-58 (**Figure 3C**).

**Figure 3.**
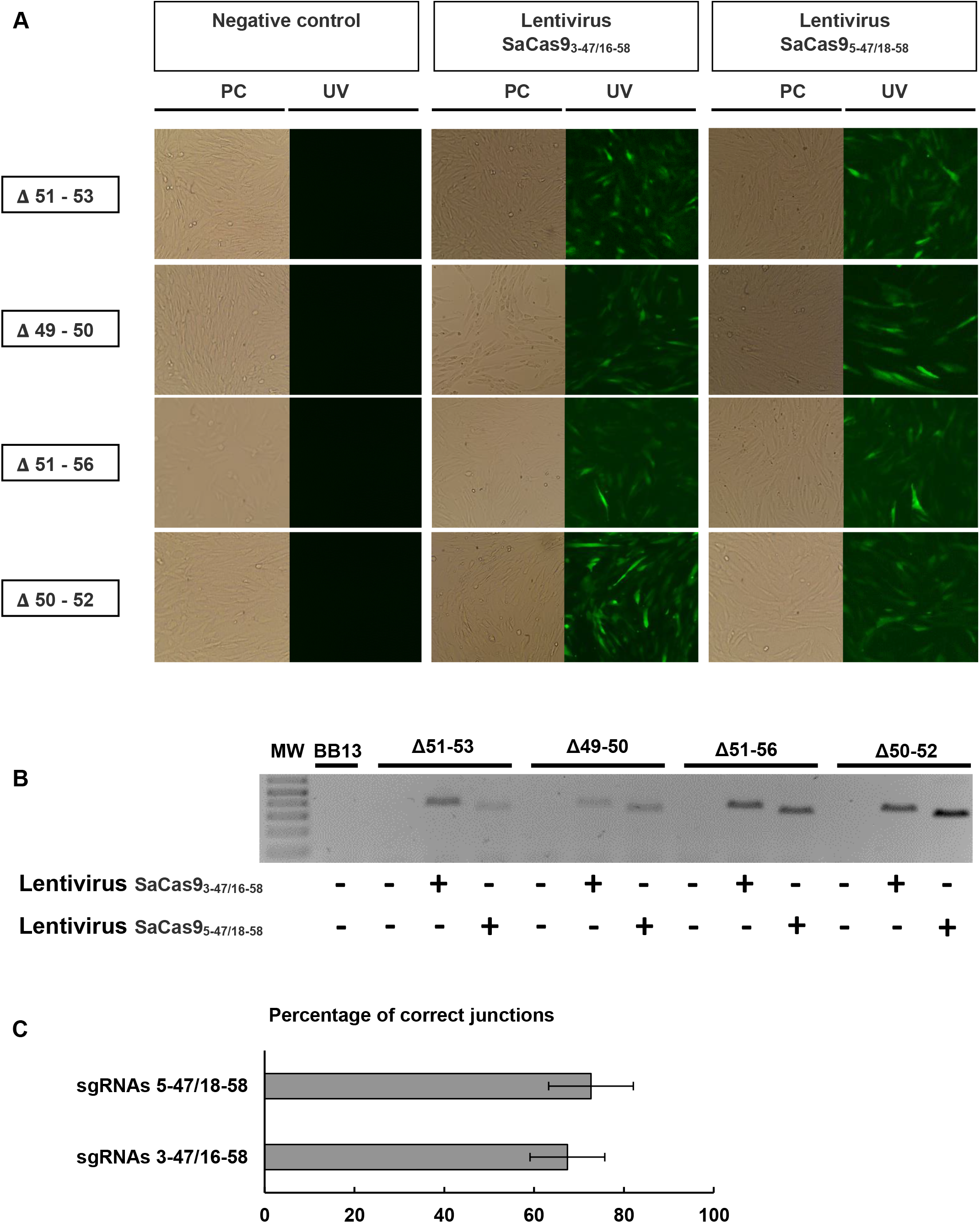
Lentiviral delivery of the SaCas9 gene and pairs of sgRNAs restored the expression of the dystrophin in DMD patient cells. Myoblasts from 4 DMD patients with different exon deletions (i.e., exons 51 to 53, exons 49 and 50, exons 51 to 56 and exons 50 to 52) were infected in 8 μg/mL of Polybrene in MB-1 medium with a third-generation lentiviral vector encoding the SaCas9 gene and a pair of sgRNAs (i.e., either the pair of sgRNAs 3-47/16-58 or the pair of sgRNAs 5-47/18-58). (A) The SaCas9 protein was immediately followed by a T2A-eGFP fragment, which fluorescence allowed to control the level of infection of the myoblasts. (B) PCR amplifications were performed from purified genomic DNA to detect the formation of the hybrid exons 47-58. (C) The amplicon sequences were submitted to TIDE analysis to determine the percentage of exact junction (n=4).

Following the confirmation of the formation of the hybrid exon, myoblasts were grown until 100% confluence was reached and allowed to fuse into myotubes for at least 7 days in low serum medium. We observed that transduced myoblasts were still able to fuse into myotubes as eGFP expression was detected in myotubes (**Figure 4A**). Next, the restoration of dystrophin expression was investigated with a western blot using an anti-dystrophin antibody (NCL-Dys2, Novocastra) in treated and non-treated myotubes (**Figure 4B**). As expected, our positive control myoblasts (derived from a healthy child) demonstrated the expression of the wild-type dystrophin (427 kDa), while no dystrophin was detected in all the myotubes from untreated DMD myoblasts. Interestingly, all myotubes generated from DMD patient myoblasts transduced with Lenti-SaCas9-3-47/16-58 or with Lenti-SaCas9-5-47/18-58 were able to express a truncated dystrophin, which size corresponded to the targeted formation of the hybrid exon 47–58 that resulted in a ~360 kDa dystrophin protein. These results demonstrated that it is possible to create a large genomic deletion that reframe the *DMD* gene and allow the *de novo* synthesis of dystrophin.

**Figure 4.**
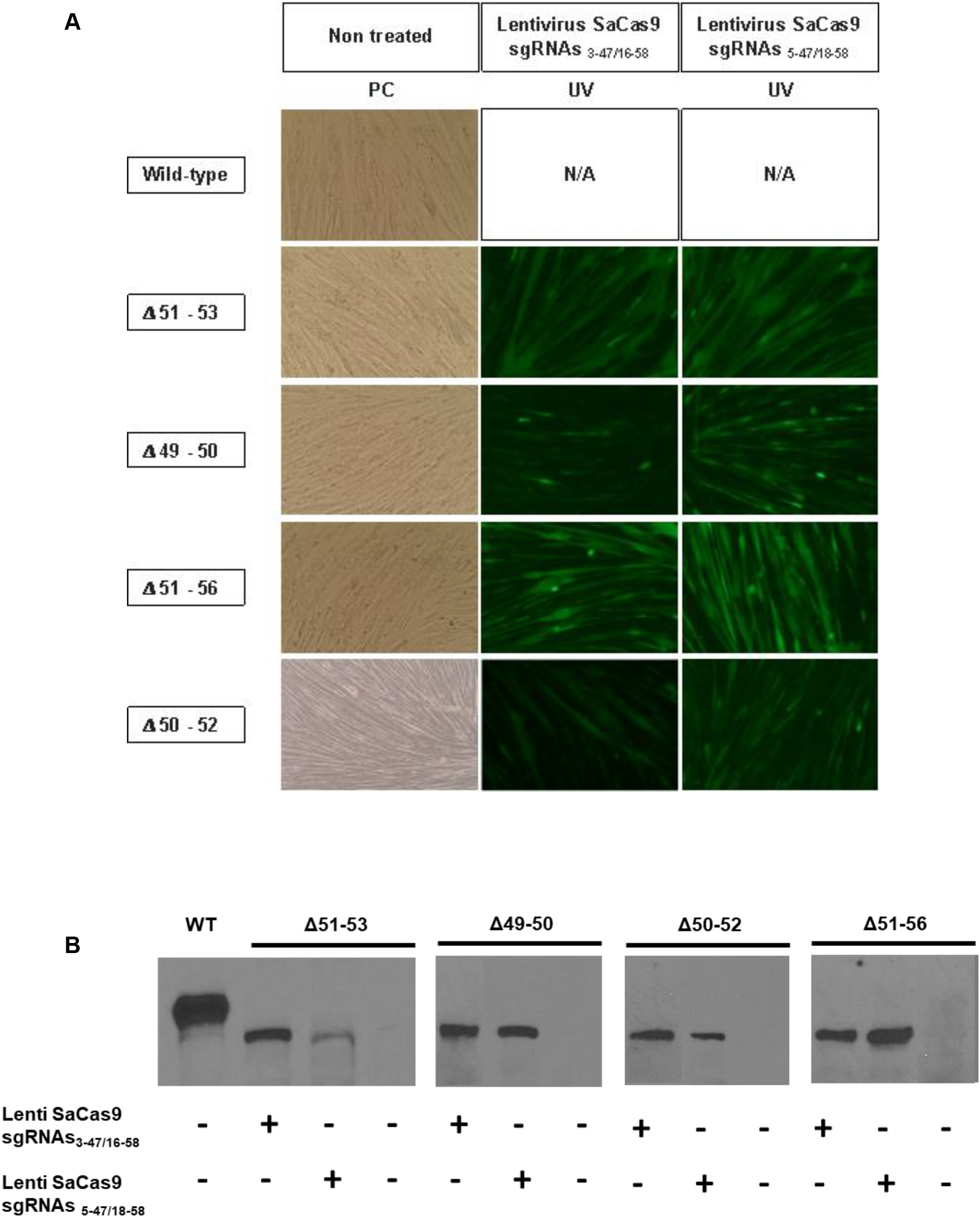
Restoration of dystrophin expression in edited myotubes. (A) Myoblasts from a healthy subject (WT) and from 4 DMD patients with different exon deletion were infected with the lentivirus coding for SaCas9, a pair of sgRNAs and eGPF, and allowed to fuse into myotubes for at least 7 days in low serum medium. EGFP expression allowed to observe that myoblasts infected with the lentivirus were able to fuse into myotubes. (B) The myotube proteins were then extracted and submitted to a western blot analysis for the detection of the dystrophin protein using an anti-dystrophin monoclonal antibody (NCL-Dys2, Novocastra).

### Off-target analysis

For the analysis of the possible off-target mutations, we focused our study on the best two pairs of sgRNAs (i.e., sgRNAs 3-47/16-58 and sgRNAs 5-47/18-58). The potential off-target sites were identified with the in silico tools provided on the website http://www.benchling.com which is calculated using an algorithm developed by Hsu et al.^19^. For each sgRNA, we considered the top 10 high scores of off-targets with the most permissive PAM, meaning NNGRR in addition to NNGRRT. We choose to focus only on off-targets related to annotated genes. For the sgRNA 3-47, the highest off-target score identified was 2,5. However, none of the 10 off-targets with the highest scores were related to an annotated gene. The sgRNA 5-47, had potential off-targets with scores below 0,8 and no annotated genes. The sgRNA 16-58 showed off-target scores ranging from 7,4 to 0,8. Its highest off-target was the IL-17A gene localized in the chromosome 6 (position +52190188). Interestingly, the off-target site in the IL-17A gene is localized in its intron 3 after the stop codon. Thus, even if off-target mutations would occur in this localization, they would have no impact on the IL-17A protein, implicated in the immune function. In addition, the sgRNA 18-58 had a potential off-target site in the TRIM67 gene localized in chromosome 1 (position - 231217983), with a score of 1,8. However, the targeted sequence is located after the STOP codon of the TRIM67 gene and thus would also have to impact on the encoded protein.

To characterize these possible off-targets, we worked with the genomic DNA of non-treated myoblasts and of myoblasts transduced with lentiviral vectors that coded for the expression of the SaCas9 and for a pair of sgRNAs. The cells were cultured for 4 weeks. IL-17A and TRIM67 off-target genomic regions were amplified by PCR and submitted to a Surveyor assay (**Supplementary data - Figure S4.A**). No difference between untreated and treated myoblasts was observed; meaning that if off-target mutations occurred, their frequency is below the detection limit of the Surveyor assay, which is 3 to 5% according to manufacturer’s instruction. The PCR products were also submitted to TIDE analysis (**Supplementary data - Figure S4.B**). Despite a slightly higher INDELs rate with the sgRNA 16-58 in IL-17A gene, no significant difference was observed in comparison to the control cells (n=4, Student T test, p = 0.34). In addition, the sgRNA 18-58 did not significantly increased INDELs rate in the TRIM67 gene (n=4, Student T test, p= 0.38).

For a deeper characterization, these possible off-target regions were submitted to deep-sequencing analysis (**Supplementary data - Figure S4.C**). For the off-target TRIM67, no significant increase of percentage of INDELs was detected between treated and untreated myoblasts (n=4, paired T-test, p = 0,6). However, the sequences of the off-target sites in the IL-17A gene exhibited a low but significant increase of INDELs at the site targeted by the sgRNA 16-58 (n=4, paired T test, p = 0, 018). The physiological impacts of these off-target mutations are probably not important since this off-target sequence is not located in an exon coding for the IL-17A protein nor in a splice acceptor or a splice donor. Consequently, no physiological effect should be produced by mutations of this off-target site.

Obviously, for any further experiment in human DMD patients, one should consider to more extensively characterize off-target events using high throughput next generation sequencing, which should be preferentially evaluated in cells of each patient because of the genomic variations from one human to another.

### Systemic delivery of AAV9 partially restores dystrophin expression in the heart of del52hDMD/mdx mice

To assess the *in vivo* feasibility of our approach, the systemic delivery of the CRISPR/Cas9 components was tested with AAV9 vectors in del52hDMD/mđx mice. Six weeks old mice were injected with a total amount of 7.5 × 10^13^ v.g./kg; 3.5 × 10^13^ v.g./kg of AAV9-SaCas9 coding for the SaCas9 gene combined with 3.5 × 10^13^ v.g./kg of AAV9-sgRNA3-47/16-58 or AAV9-sgRNA5-47/18-58 coding for the production of a pair of sgRNAs that previously permitted the formation of a hybrid exon 47-58. Mice were sacrificed 6 weeks after the injection to ensure the accumulation of dystrophin protein in edited cells. In saline injected del52hDMD/mđx mice, only a few dystrophin positive cardiomyocytes were observed by immunohistochemistry. Since our polyclonal antibody rabbit anti-dystrophin detected mouse and human dystrophin, the dystrophin positive cardiomyocytes may be due to somatic mutations of the human or mouse DMD gene restoring the normal reading frame. This is similar to the so called revertant skeletal muscle fibers observed in the *mdx* mice^20^ In all mice treated with a pair of sgRNAs, either sgRNAs 3-47/16-58 or sgRNAs 5-47/18-58, dystrophin expression was observed in more cardiomyocytes. Interestingly, a higher dystrophin expression level was observed by immunohistochemistry when the sgRNAs 3-47/16-58 was used (**Figure 5A**). The restoration of dystrophin expression was consistent with the detection of the hybrid exons 47-58 by PCR (**Figure 5B**). However, such PCR did not permit the detection of hybrid exons neither in the *Tibialis anterior* nor in the diaphragm of the treated del52hDMD/mdx mice. This may be due to a lower amount of viral particles in these tissues (data not shown). TIDE analysis demonstrated that the hybrid exons generated by the sgRNAs 3-47/16-58 harboured the expected sequence in 86 % of amplicons and that those produced following treatment with sgRNA 5-47/18-58 contained 83 % of the right nucleotide sequence (**Figure 5D**). Following protein extraction, the western blot analysis confirmed that the *de novo* expressed dystrophin in the heart, corresponded with the size of the internally truncated dystrophin that we aimed to create (**Figure 5C**). Together, these results demonstrate that our approach is suitable for *in vivo* applications.

**Figure 5.**
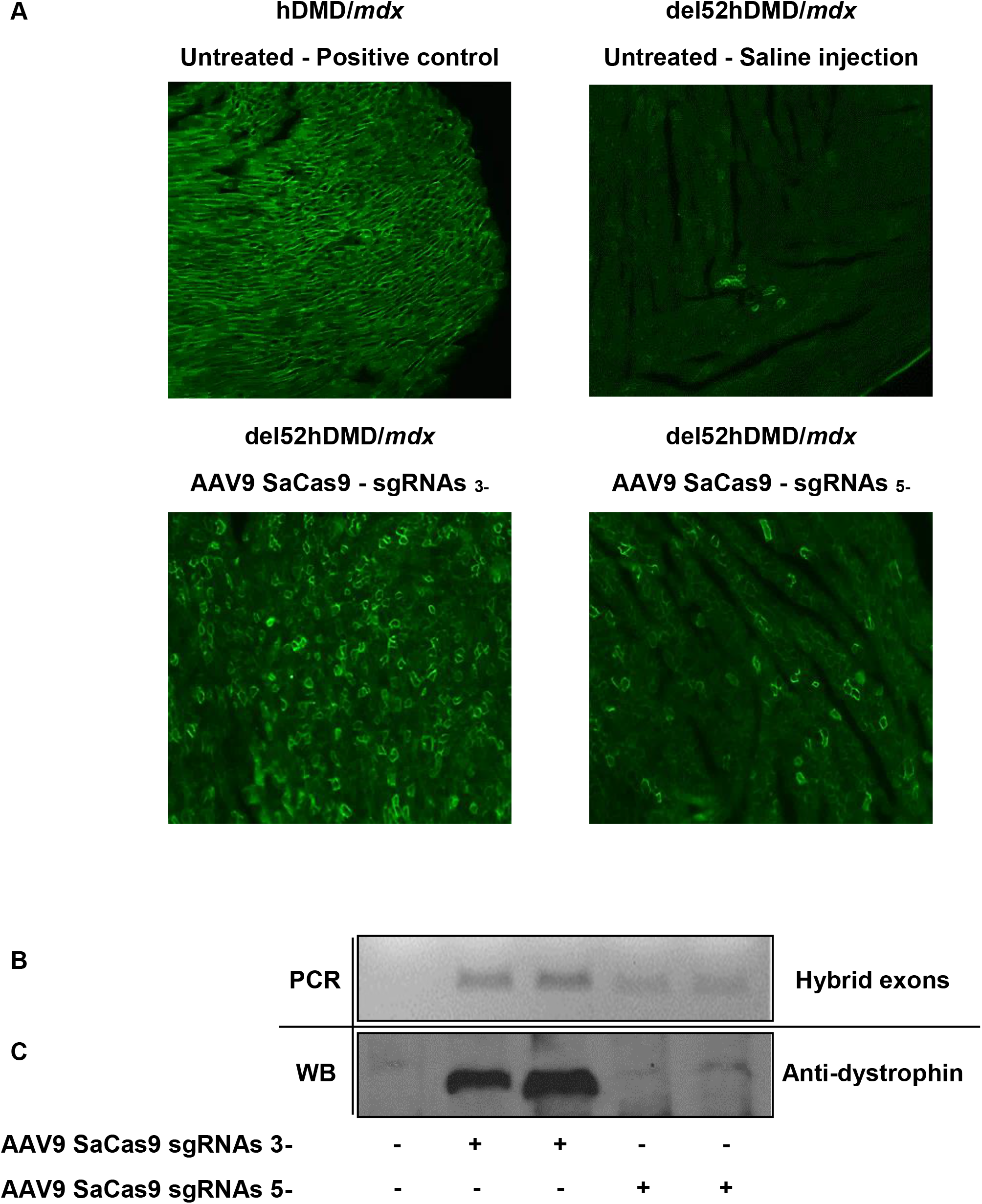
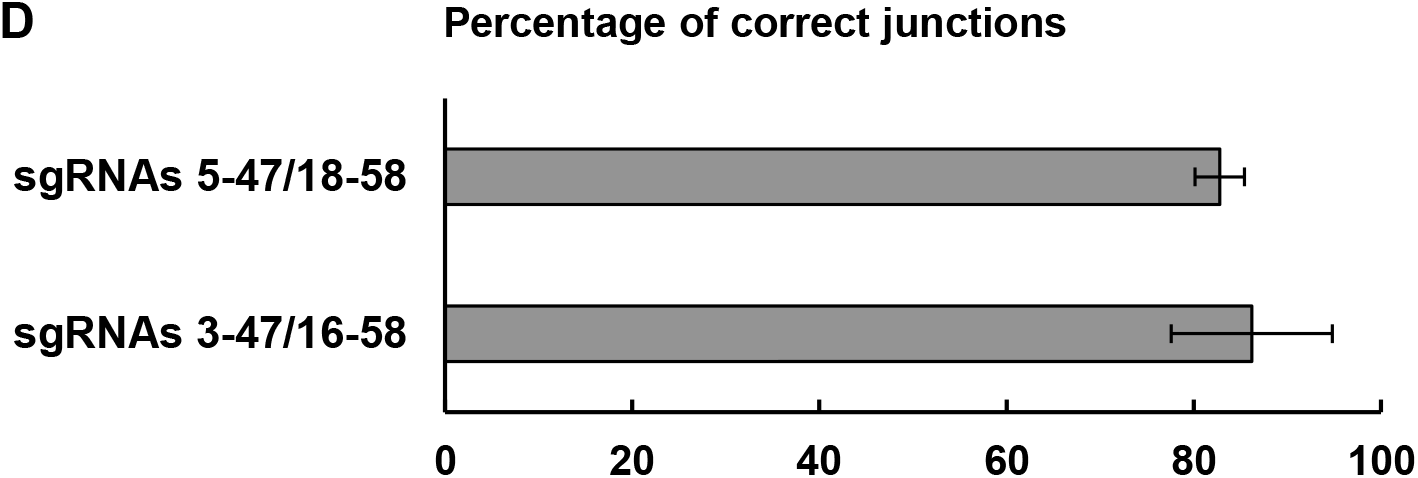
Formation of the hybrid exons 3-47/16-58 and 5-47/18-58 restored the expression of dystrophin protein *in vivo* in cardiomyocytes of the del52hDMD/mđx mouse model. 4-5 weeks old del52hDMD/mdx were systemically injected in the tail vein with a total amount of 1,5.10^12^ v.g. of AAV9s coding for the SaCas9 and for a pair of sgRNAs (i.e., either sgRNAs 3-47/16-58 or sgRNAs 5-47/18-58). Mice were euthanized 6 weeks after the injection and tissues were collected. (A) Heart cryo-sections were stained with a rabbit polyclonal anti-dystrophin antibody (ab15277, Abcam). (B) PCR confirmed the formation of the hybrid exons 47-58 in the heart of treated mice. (C) The presence of the internally truncated dystrophin protein was confirmed by western blot analysis. (D) The amplicon sequences of the hybrid exons were submitted to TIDE analysis to determine the percentage of exact junction (n=4).

## DISCUSSION

Over the past five years, several CRISPR-based approaches have been investigated for the development of therapies for DMD. Single-cut genome editing relies on the alteration of splice-acceptor sequences to splice out the targeted exon to restore the normal reading frame in the mature mRNA leading to the production of dystrophin protein^21–26^ Other groups develop strategies that rely on the complete removal of one or several exons to restore the *DMD* gene reading frame using a pair of sgRNAs inducing DSB in introns^26–28^. The edition of the dystrophin gene by Homology Directed Repair (HDR) has also been investigated, by the treatment of muscle stem cells followed by their engraftment or by the direct delivery of nanoparticles formed by a complex between a Cas9/sgRNA and a donor DNA^26, 29,30^. This approach restored the dystrophin gene and full-length dystrophin protein, but it remains limited to *in situ* treatment only.

Current strategies, that rely on exon skipping or deletion of one or several complete exons, do not consider the structure of the dystrophin protein and the phasing of the SLR that results from splicing out or deletions of these exons. As reported by Le Rumeur’s group^6^ the abnormal structure of the SLRs that form the central rod domain of the dystrophin protein could be responsible for the severity and disease progression of Becker Muscular Dystrophy (BMD). Indeed, dystrophin from some BMD patients harbouring large in-frame deletions can still be functional and lead to a milder phenotype^31^. Thus, features learned from BMD patients is of major interest for research groups that develop mini and micro-dystrophin whose works mainly focus on the designing a truncated dystrophin protein, which nevertheless contained adequate functional structural domain^32^ Interestingly, it has been shown that microdystrophins having SLRs correctly phased improved its function in comparison to abnormal SLRs.

Several researchers have worked for many years to produce a mini or a microdystrophin gene that not only produces a protein but that also codes for a protein with an optimal structure to insure a good function^33–40^. Since an adequate structure is a very important for a mini and a micro-dystrophin gene, the structure of the dystrophin protein produced by a CRISPR/Cas9 modification should also be carefully taken in consideration. In a previous work, we used the SpCas9 to induce precise deletion to form the hybrid exon 50-54, through induction of double strand breaks at precise genomic sites. The main advantage of the SaCas9 over the SpCas9 is that its smaller size could in principle permit to insert its gene as well as 2 sgRNAs in only one AAV. This great advantage, for an eventual clinical application, would not only reduce the cost of the treatment but moreover every cell infected by the unique AAV vector would have all the components required to produce the hybrid exon. However, the development of such product requires further development to obtain this all in one vector, which requires a promoter smaller than the CMV that allow expression of SaCas9 in skeletal muscle and cardia cells only, along with sgRNA promoter that exhibits enough activity.

Consequently, our research group considered that it could be highly relevant to focus on the development of a CRISPR-based therapy that not only restores the *DMD* gene reading frame but also aims to form a normal SLR that mimics the dystrophin protein present in BMD patients with no severe symptoms. The aim of our project was to provide proof of principle for a therapeutic approach using the CRISPR/SaCas9 technology for DMD patients harbouring different mutations in their *DMD* gene. In a previous publication, our team demonstrated that through the use of the Cas9 of *Streptococcus pyogenes* (SpCas9) it was possible to form a hybrid exon 50-54 that not only restored the *DMD* gene normal reading frame but also formed a normally phased SLR, a key component of the central rod domain of the dystrophin protein. However, this strategy was only suitable for DMD patients affected by a mutation of their *DMD* gene between exons 50 and 54. Based on a similar approach, we aimed in the present project to form a hybrid exon that could provide a therapy for a higher percentage of DMD patients. In addition, we choose to design our approach using SaCas9, which is smaller than SpCas9 and could thus be delivered by a single AAV coding for the SaCas9 gene and a pair of sgRNAs.

Previous works had already demonstrated the ability of the SaCas9 nuclease and a pair of sgRNAs to edit the *mdx* gene in the *mdx -/-* mouse, a dystrophic model, by removing completely the mutated exon 23 that consequently restored the expression of dystrophin^26, 27,41^. Here, we report for the first time the use of the SaCas9 to edit the human *DMD* gene in myoblasts from DMD patients and in the humanized dystrophic del52hDMD/mdx mouse model. We initially aimed to identify pairs of sgRNAs that could be suitable for the treatment of DMD patients having a deletion of exon 50. Finally, we identified two pairs of sgRNAs (sgRNAs 3-47/16-58 and sgRNAs5-47/18-58) that could be used in a therapy for all DMD patients having a deletion, an insertion or a point mutation between exon 47 and 58. Note that we purposely did not select sgRNA pairs that would delete exons 44 or 45, which are coding for dystrophin segment binding with nNOS, since this interaction is important for a normal dystrophin function and muscle fibres integrity^42, 43^. Since DMD is responsible of muscle wasting development of fibrosis that reduces the amount of targetable muscle tissue and thus lead to impaired muscle contraction, this therapeutic approach should be primarily used for the treatment of young DMD patients. On the other hand, our selected pairs of sgRNAs could also be used to treat BMD patients who exhibit a severe phenotype and thus could be of interest for the structural improvement of the dystrophin of Becker patient with severe phenotype. The potential of this strategy should be explored, but to date no mouse model harbouring severe Becker phenotype exists and thus this precludes the study of the benefits of the improvement of the structure of the dystrophin protein in such patient.

The pair of sgRNAs 3-47/16-58 seems the most promising as it allowed the restoration of the dystrophin protein expression more efficiently than the sgRNA 5-47/18-58 pair. The sgRNA 16-58 could lead to an off-target mutation in the IL-17A gene, which could be problematic in DMD patients. Even if this off-target site is located within a non-coding segment of the IL-17A gene, strategies to limit possible adverse effects would have to be considered. A possible strategy would be to use a skeletal muscle and cardiac tissue specific promoter, such as the CK8e promoter developed by Hauschka’s laboratory^21, 26,44^, which restricts the expression of the nuclease to tissues where editing is required to improve the DMD patient phenotype. In addition, the development of tools, which prevent the sustained expression of CRISPR nucleases, are required to restrain the nuclease expression to the minimal time period required for a sufficient editing of the target gene^45, 46^, while limiting the occurrence of off-target mutations. Here we empirically assessed the potential off-targets of our sgRNAs based on an in silico prediction tool. This tool, and other similar ones, identify off-targets based on the position of one or several mismatches in the sgRNA^19, 47^ but several articles have reported that the sites of predicted off-target mutations are not always reliable^48^. Ultimately, a pre-clinical study should characterize more extensively the overall off-target mutations of sgRNA 3-47 and 16-58 using un-biased high throughput next generation sequencing^49–51^.

Our follow-up study will aim to improve the overall *DMD* gene editing *in vivo*, in young del52hDMD/mdx mouse, to restore dystrophin protein expression in the diaphragm and skeletal muscle in addition to the heart. We will also assess whether the formation of hybrid exons, such as the hybrid exon 47-58, which maintains the SLR structure, results in a more substantial functional improvement in comparison to the editing of the *DMD* gene by the removing of one or several full exons, without maintaining the correct phasing of the SLR.

## MATERIALS & METHODS

### Identifications of DNA targets and sgRNA cloning

For the identification of the sgRNA targets, we used the website www.benchling.com. Based on the *DMD* gene sequence (ENSG00000198947), we identified all the PAMs “NNGRRT” for the Cas9 for *Staphylococcus aureus* from in the exons 46 to 58. Oligonucleotides corresponding to the sgRNAs of interest were ordered from Integrated DNA Technology (Coralville, IA).

### Expression vector and cloning of sgRNAs

Selected sgRNAs were cloned into the plasmid pX601-AAV-CMV::NLS-SaCas9-NLS-3xHA-bGHpA;U6::BsaI-sgRNA (PX601) (Addgene #61591) following Zhang’s laboratory protocol. The plasmid PX601 was linearized using the BsaI restriction enzyme (NEB, Ipwisch, MA) followed by purification with a gel extraction kit (Thermo Scientific, Vilnius, Lithuania). The sgRNAs of interest were then cloned into the BsaI site using the Quickligase (NEB, Ipwisch, MA) followed by transformation in Top10 competent bacteria. In the plasmid pBSU6_FE_ScaffoldRSV_GFP (gift from Dirk Grimm Lab, Heidelberg University, Germany), oligonucleotides were sequentially cloned into the BsmbI or BbsI restriction sites. Clones were amplified in liquid LB and following plasmid DNA extraction, clones were sequenced. Sequencing results were analysed using CLC Sequence Viewer (CLC Bio).

### Transfection procedure

293T cells were grown under 5% CO2 at 37°C with DMEM supplemented with 10% FBS and 1% penicillin/streptomycin. The initial experiments to assess the activity of sgRNAs were made in 293T cells. A total amount of 900 ng of plasmid was transfected into cells using 3 μL of Lipofectamine 2000. 48 h following transfection, cells were collected and genomic DNA was extracted and purified using the phenol: chloroform method and concentrated by a salt/ethanol precipitation.

### Genomic DNA extraction and analysis

48 hours after transfection of the pX601-AAV-CMV:NLS-SaCas9-NLS-3xHA-bGHpA;U6::BsaI-sgRNA plasmid(s), the genomic DNA was extracted from the 293T cells or myoblasts using a standard phenol: chloroform method. Briefly, the cell pellet was suspended in 100 μl of lysis buffer containing 10% sarcosyl and 0.5 M pH 8 ethylenediaminetetraacetic (EDTA). Twenty (20) μl of proteinase K (10 mg/ml) were added. The suspension was mixed by up down and incubated 10-15 min at 55°C. Suspension was then centrifuged at 13200 rpm for 5 min. The supernatant was collected in a new microfuge tube. One volume of phenol-chloroform was added and following centrifugation, the aqueous phase was recovered in a new microfuge tube. Then DNA was precipitated using 1/10 volume of NaCl 5 M and two volumes of 100% ethanol followed by 5 min centrifugation at 13200rpm. The pellet was washed with 70% ethanol, centrifuged and the DNA was suspended in double-distilled water. The genomic DNA concentration was assayed with a Nanodrop^TM^ spectrophotometer (Thermo Scientific, Logan, UT).

### Assessment of the formation by sgRNAs of INDELs on on-target and off-target

Genomic region targeted by a sgRNA (either on-target or off-target sequences) were amplified by PCR before further analysis. First, cleavage efficiency was assessed by Surveyor assay according to manufacturer’s instruction. PCR products were then sequenced and the resulting.ab files were used for their characterization using the TIDE tool provided on https://tide.deskgen.com^52^. Amplifications of genomic DNA from untreated cells were used as control sample. For deep-sequencing analysis, new sets of primers were designed to permit the amplification of PCR products no longer than 350 bp.

### Assessment of the formation of hybrid exons

For the characterization of the nucleotide sequence of the hybrid exons, PCR product corresponding to the amplification of these hybrid exons were cloned into pMiniT plasmid vector of a PCR cloning kit following manufacturer’s instruction (NEB, Ipswich, USA). After transformation into bacteria and clone growth in a liquid medium, plasmidic DNA was extracted and purified for sequencing.

The TIDE analysis tool was used to characterize of the rate of formation of perfect hybrid exons. As a control sample, hybrid exons that contained the exact nucleotide sequence, previously inserted in a pMiniT vector and sequenced, were amplified by PCR and were used as control sample for the analysis. As a test sample, pools of PCR products amplified from treated cells were used.

### Lentivirus production

A lentiviral vector that allows for the expression of the SaCas9 along with 2 sgRNAs was constructed starting from the plasmid FUGW (Addgene # 14883). First, the SaCas9 gene and a T2A-eGFP were inserted under the control of the CMV promoter. Two U6 promoters were then inserted to control the expression of two sgRNAs. A restriction site, either BbsI or BsmbI, was inserted downstream of each U6 promoter to permit the cloning of each individual sgRNA. For each lentivirus production, 4.10^6^ 293T (packaging cell line) were plated into a 10×10 cm petri-dish the day prior transfection. The day of transfection, cells were transfected with a combination of four plasmids; Gag-Pol, Rev, VSV-G and the transfer vector, using a Calcium Phosphate transfection method. The day after the transfection, medium was removed and replaced with fresh medium. 48 hours later, medium containing the viral particles was collected. Following centrifugation to remove cell debris, viral supernatant was concentrated and purified by ultracentrifugation at 15,000 g for 90 min in 20% sucrose gradient. Following supernatant removal, viral particles were suspended into cold PBS 1X.

### Myoblast transduction with recombinant lentiviral vector

Myoblasts from different DMD patients (harbouring a deletion of exons 49 to 50, 50 to 52, 51 to 53, or 51 to 56) were grown under 5% CO2 at 37°C in MB-1 medium supplemented with 40 ng/mL of bFGF and 15% of FBS. For viral transduction, myoblasts were treated overnight in MB-1 medium supplemented with 8 μg/mL of Polybrene with the previously purified lentivirus. The day after the transduction, cells were washed using PBS 1X and cultured in Polybrene free MB-1 medium. Transduction efficiency was assessed 48 h after medium replacement by observing eGFP fluorescence.

### Western blot analysis

After the fusion of myoblasts into myotubes, proteins were extracted using a protein lysis buffer (75 mM Tris-HCl pH 8.0, 1 mM DTT, 1 mM PMSF, 1% SDS). The protein concentrations were determined using an amido-black assay^53^. Protein samples were separated onto a 6% SDS-polyacrylamide electrophoresis gel. Proteins were then transferred overnight onto a nitrocellulose blotting membrane. That membrane was blocked for 8 hours in 5% milk in TBS 1X and 0.01% Tween-20 at 4 °C. The primary antibody anti-dystrophin (dilution 1:50, NCL-Dys2, Novocastra, Newcastle, UK) was incubated overnight at 4°C. The secondary antibody goat anti-mouse HRP (dilution 1:5000) was applied for 1 h. Blots were revealed using the Clarity™ Western ECL Blotting Substrates (Bio-Rad Laboratories, Mississauga, ON, Canada).

### Production of AAV9 viral vectors

All viral vectors were produced by the Molecular Tools Platform of the CRIUSMQ (Québec, Canada). The AAV9 vector that encodes the SaCas9 gene under the control of the CMV promoter was produced from the plasmid PX601 (Addgene Inc., Cambridge, MA). Two AAV9 vectors that permitted the expression of a pair of sgRNAs were produced from the plasmid pBSU6 (pBSU6_FE_Scaffold_RSV_GFP, Dirk Grimm Lab, Heidelberg University, Germany).

### *In vivo* experiments

Three AAV9 vectors were designed to determine the *in vivo* feasibility of the formation of the hybrid exons 47-58. One AAV9 permitted the expression of the SaCas9 protein under the control of the CMV promoter. The two other AAV9s allowed the production of two pairs of sgRNAs: one vector encoded the sgRNA 3-47 (targeting the exon 47) and the sgRNA 16-58 (targeting the exon 58), and the other vector encoded the sgRNA 5-47 (targeting the exon 47) and the sgRNA 18-58 (targeting the exon 58). The sgRNA expression was controlled by a pU6 human promoter. Virus that coded for the pairs of sgRNAs also coded for the expression of the Green Fluorescent Protein (GFP). Six weeks after the intravenous injection, mice were sacrificed and tissues (heart, diaphragm, *Tibialis anterior*, liver) were collected and incubated overnight in 30% sucrose at 4°C. Then, samples were embedded into cryomatrix and flash frozen. Samples were stored at −80°C.

### Targeted deep-sequencing and bioinformatics analysis

For deep-sequencing analysis, primers were redesigned to allow the amplification of PCR products of a maximum of 350 bp. Samples were sent for sequencing to the Next-Generation Sequencing platform of McGill University (McGill University and Genome Quebec Innovation Centre, Montreal, Canada). Fastq files quality were checked using the FastQC tool (https://www.bioinformatics.babraham.ac.uk/projects/fastqc/). Before characterization of sequencing results, Fastq files were trimmed by sliding window (SLIDINGWINDOW:4:15) and minimum length (according to the expected size of the amplicons) using the algorithm Trimmomatic^54^.

### Molecular modelling

Homology models of WT *DMD* gene corresponding to spectrin-like repeats R18 and R23 were obtained from the eDystrophin web site (http://edystrophin.genouest.org/)^18^. Primary sequence alignment was realized using the TM-Align^55^ (http://zhanglab.ccmb.med.umich.edu/TM-align/) web server based on the structural alignment of R19 and R21 homology models. Homology models were realized using the iTasser web server (http://zhanglab.ccmb.med.umich.edu/I-TASSER/)^56–59^ using primary sequences of hybrid exon for deletion of exons 47-58.

### HDMD mouse model

The hDMD/mdx mouse (gift from Dr. A Aartsma-Rus) contains the full human *DMD* gene (2.4 Mb) (coding for dystrophin) stably integrated into the mouse chromosome 5^60^. In addition, the mouse dystrophin gene *(mdx* gene) contains a point mutation in the exon 23, which creates a stop codon thus abrogating the production of mouse dystrophin. The del52hDMD/mdx (gift from Dr. A. Aartsma Rus) also contains the human *DMD* gene but harbours a deletion of the exon 52 that abrogates the synthesis of the human dystrophin protein^61^. All experiments were approved by the animal care committee of the Centre Hospitalier de l’Université Laval.

### Immunohistochemistry

Frozen tissues were processed with a Cryostat (Leica Biosystem Inc., Concordia, ON, Canada) to obtain 12 μm thick transversal sections that were placed on a glass-slide pre-coated with glycine. Tissue sections were blocked for 1 h in a blocking solution (PBS 1X supplemented with 10% of FBS. Primary antibody directed against dystrophin (ab15277, Abcam Inc., Toronto, ON, Canada), diluted in the blocking solution was incubated for 1 h. After 3 washing steps of 10 min with PBS 1X, a secondary antibody, i.e., a goat anti-rabbit IgG Alexa Fluor 488 (A-11008, RRID AB_143165, Thermo Fisher Scientific), diluted in the blocking solution, was then added for another 1 h. Next, samples were wash 3 times for 10 min with PBS 1X, then stained samples were mounted under a covered glass-slide with PBS 1X supplemented with 50% of glycerol. Dystrophin expression and localization was observed by fluorescence microscopy.

## Acknowledgments

We thank Dr Aartmus for providing us with the del52hDMD/mdx mouse model. This research project was supported by grants from Canadian Institute of Health Research, Foundation for Cell and Gene Therapy and TheCell Network. Duchêne has been supported by a fellowship from MITACs.

The authors declared no competing financial interests.

## Author Contributions

Duchêne designed experiments, did the experiments, and wrote the manuscript. Cherif technically assisted for western blots. Lyombe-Engembe assisted for the designing of the experiments. Rousseau technically assisted for molecular biology. Ouellet technically assisted for lentiviral vector production. Lagüe and Barbeau made the model of the structure of spectrin-like repeats. Tremblay conceived the experiments and corrected the manuscript.

